# Evaluation of efficacy- *versus* affinity-driven agonism with biased GLP-1R ligands P5 and exendin-F1

**DOI:** 10.1101/2021.04.01.438033

**Authors:** Amaara Marzook, Shiqian Chen, Phil Pickford, Maria Lucey, Yifan Wang, Ivan R Corrêa, Johannes Broichhagen, David J Hodson, Victoria Salem, Guy A Rutter, Tricia M Tan, Stephen R Bloom, Alejandra Tomas, Ben Jones

**Affiliations:** Section of Endocrinology and Investigative Medicine, Imperial College London, London, United Kingdom; Section of Cell Biology and Functional Genomics, Imperial College London, London, United Kingdom; New England Biolabs, Ipswich, MA, USA; Leibniz-Forschungsinstitut für Molekulare Pharmakologie, Berlin, Germany; Institute of Metabolism and Systems Research (IMSR), University of Birmingham, Birmingham, United Kingdom; Centre for Endocrinology, Diabetes and Metabolism, Birmingham Health Partners, Birmingham, United Kingdom; Centre of Membrane Proteins and Receptors (COMPARE), Universities of Birmingham and Nottingham, Midlands, United Kingdom; Department of Bioengineering, Imperial College London, London, United Kingdom; Lee Kong Chian School of Medicine, Nanyang Technological University, Singapore

**Keywords:** GLP-1R, biased agonism, endocytosis, exendin-4, β-arrestin

## Abstract

The glucagon-like peptide-1 receptor (GLP-1R) is an important regulator of glucose homeostasis and has been successfully targeted for the treatment of type 2 diabetes. Recently described biased GLP-1R agonists with selective reductions in β-arrestin *versus* G protein coupling show improved metabolic actions *in vivo*. However, two prototypical G protein-favouring GLP-1R agonists, P5 and exendin-F1, are reported to show divergent effects on insulin secretion. In this study we aimed to resolve this discrepancy by performing a side-by-side characterisation of these two ligands across a variety of *in vitro* and *in vivo* assays. Exendin-F1 showed reduced acute efficacy *versus* P5 for several readouts, including recruitment of mini-G proteins, G protein-coupled receptor kinases (GRKs) and β-arrestin-2. Maximal responses were also lower for both GLP-1R internalisation and the presence of active GLP-1R-mini-G_s_ complexes in early endosomes with exendin-F1 treatment. In contrast, prolonged insulin secretion *in vitro* and sustained anti-hyperglycaemic efficacy in mice were both greater with exendin-F1 than with P5. We conclude that the particularly low acute efficacy of exendin-F1 and associated reductions in GLP-1R downregulation appear to be more important than preservation of endosomal signalling to allow sustained insulin secretion responses. This has implications for the ongoing development of affinity- *versus* efficacy-driven biased GLP-1R agonists as treatments for metabolic disease.

## 1 Introduction

With the increasing worldwide prevalence of type 2 diabetes (T2D) [1], there is an urgent need for more effective drugs to treat this condition. T2D results from a combination of relative insulin deficiency and resistance to its action in central and peripheral tissues, and is commonly associated with excess adiposity or obesity [2]. The glucagon-like peptide-1 receptor (GLP-1R), a class B G protein-coupled receptor (GPCR) expressed at high levels in pancreatic beta cells and at lower levels in anorectic centres in the brain, is a well-established target for pharmacological T2D treatment [3]. GLP-1R activation augments glucose-stimulated insulin secretion, improves beta cell survival and suppresses appetite, with the latter resulting in weight loss and improvements in insulin sensitivity [4]. Pharmacokinetically optimised GLP-1R agonists based on the amino acid sequence of either the cognate agonist GLP-1(7-36)NH_2_ or its homologue exendin-4 [5] are approved for the treatment of T2D. These agents not only improve glycaemic control and induce weight loss but also reduce cardiovascular [6] and all-cause mortality [7] in people with T2D.

Recent preclinical studies have shown that GLP-1R agonists that favour G protein signalling and generation of cyclic adenosine monophosphate (cAMP) over β-arrestin recruitment are particularly effective at reducing blood glucose levels [8–12]. Moreover, it is suggested that the markedly reduced β-arrestin recruitment seen at the GLP-1R with the dual GLP-1R/glucose-dependent insulinotropic peptide receptor (GIPR) agonist Tirzepatide [13–15] may contribute to its superior anti-diabetic efficacy in clinical trials [16]. An appealing explanation for the effects of these “biased” GLP-1R agonists is that reductions in β-arrestin-mediated desensitisation, as well as a trafficking phenotype that favours preservation of GLP-1R at the plasma membrane, lead to prolonged intracellular signalling and cumulatively greater insulin release over time. However, this has not been demonstrated for all published examples of biased GLP-1R agonists. In particular, “P5”, the first bespoke biased GLP-1R agonist to be described, was potently anti-hyperglycaemic but poorly insulinotropic *in vivo*, with its metabolic effects instead partly attributed to increases in adipogenesis [8]. This contrasts with “exendin-phe1” [9] (referred to here as “exendin-F1”), a peptide that showed marked increases in insulin release compared to exendin-4 in cellular models and in mice. One possible explanation for this discrepancy is that different approaches were used to evaluate the two ligands, with exendin-F1 tested using prolonged incubations with beta cells and islets, as well as sub-chronic *in vivo* studies, to specifically seek functional evidence of reduced desensitisation over the course of several hours [9]. However, to date, P5 has not been examined in this way.

Establishing a consensus mechanism of action for biased GLP-1R agonists would help guide their development for the treatment of T2D and related metabolic diseases. In the present work we perform direct pharmacological comparisons of P5 and exendin-F1, with exendin-4 included as the reference peptide. As well as determining relative preferences for G protein and β-arrestin recruitment responses, we focussed in particular on differences in GLP-1R membrane trafficking and activation in different endomembrane compartments [17]. Our study highlights a number of pharmacological properties that diverge between P5 and exendin-F1, suggesting these GLP-1R agonists may in fact possess distinct modes of action.

## 2 Materials and methods

### 2.1 Peptides

Exendin-4, exendin-F1 and P5 were all custom synthesised by Wuxi Apptec with >90% purity confirmed by HPLC. The synthesis of Luxendin645 and exendin-4-Cy5 have been described previously [18,19]

### 2.2 Cell culture

HEK293T cells were maintained in DMEM supplemented with 10% FBS and 1% penicillin/streptomycin. HEK293-SNAP-GLP-1R cells, generated by stable transfection of pSNAP-GLP-1R (Cisbio) into HEK293 cells [20], were maintained in DMEM supplemented with 10% FBS, 1% penicillin/streptomycin and 1 mg/ml G418. PathHunter CHO-K1-GLP-1R-βarr2-EA cells (DiscoverX) were maintained in Ham’s F12 medium with 10% FBS and 1% penicillin/streptomycin. INS-1 832/3 cells (a gift from Prof Christopher Newgard, Duke University) [21], and subclones thereof lacking endogenous GLP-1R or GIPR after deletion by CRISPR/Cas9 (a gift from Dr Jacqueline Naylor, AstraZeneca) [22], were maintained in RPMI at 11 mM glucose, supplemented with 10% FBS, 10 mM HEPES, 1 mM pyruvate, 50 μM β-mercaptoethanol, and 1% penicillin/streptomycin.

### 2.3 GLP-1R competitive binding measurements

Cells were labelled with SNAP-Lumi4-Tb (Cisbio, 40 nM, 60 min at 37°C, in complete medium). After washing to remove unbound probe, cells were resuspended in HBSS with 0.1% BSA and metabolic inhibitor cocktail (20 mM 2-deoxyglucose and 10 mM NaN_3_) to inhibit endocytosis, as previously described [9,23], and seeded into white opaque plates. After 20 min at room temperature, cells were then placed at 4°C, and a range of concentrations of test ligands were applied concurrently with 10 nM Luxendin645 [19] or 5 nM exendin-4-Cy5, with a range of concentrations of Luxendin645 or exendin-4-Cy5 also applied to establish equilibrium binding parameters for the competing labelled GLP-1R probe. After a 24-hour incubation period at 4°C, binding was measured by TR-FRET using a Spectramax i3x plate reader (Molecular Devices) fitted with an HTRF module.

### 2.4 cAMP assays

Resuspended cells were stimulated at 37°C in their respective serum-free medium in 96-well half area opaque white plates before addition of HTRF lysis buffer and detection reagents (cAMP Dynamic 2 kit, Cisbio). The duration of stimulation and inclusion or not of the phosphodiesterase inhibitor 3-isobutyl-1-methylxanthine (IBMX) is indicated in the relevant figure legend. The assay was read by HTRF.

### 2.5 Homologous desensitisation assay in beta cells

INS-1 832/3 cells were seeded into poly-D-lysine-coated 96-well plates in complete medium at 11 mM glucose in the presence of different concentrations of test agonist. After an overnight incubation, medium was removed and cells were washed 3 times in HBSS, followed by a 1-hour resensitisation period in complete medium. Cells were then stimulated at 37°C with 100 nM GLP-1 + 500 μM IBMX for 10 min followed by lysis and cAMP determination as described in Section 2.4.

### 2.6 β-arrestin-2 recruitment by enzyme complementation

PathHunter CHO-K1-GLP-1R-βarr2-EA cells were stimulated for 30 min at 37°C in serum-free Ham’s F12 medium prior to addition of lysis / detection reagents (DiscoverX). The assay was read by luminescence.

### 2.7 NanoBiT mini-Gs andβ-arrestin-2 GLP-1R recruitment assays

These assays were performed as described previously [24]. HEK293T cells were transiently transfected using Lipofectamine 2000 with 50 ng each of GLP-1R-SmBit and LgBit-β-arrestin-2 (Promega) diluted in pcDNA3.1, or with 500 ng each of GLP-1R-SmBiT and LgBiT-mini-G_s_ (a gift from Prof Nevin Lambert, Medical College of Georgia) [25], per well of a 12-well plate, and the assay was performed 24 hours later. For the kinetic mode assay, cells were resuspended in HBSS containing Furimazine (1:50), seeded into half-area opaque white plates, and total luminescent signal at baseline was recorded over 5 min at 37°C using a Flexstation 3 plate reader. Ligands were then added, and signal was serially monitored for up to 30 min. Ligand-induced change was expressed relative to baseline values for each well. For the endpoint mode assay, cells were resuspended in HBSS and seeded into half-area opaque white plates containing prepared ligands. After a 5-minute incubation at 37°C, Furimazine prepared in HBSS was added, and luminescent signal was recorded over 3 min using a Spectramax i3x plate reader.

### 2.8 NanoBRET GLP-1R recruitment assays

GRK2-Venus, GRK5-Venus and GRK6-Venus were gifts from Prof Denise Wootten, Monash University. Nb37-GFP [26] was a gift from Dr Roshanak Irannejad, University of California, San Francisco. HEK293T cells were transfected using Lipofectamine 2000 with 50 ng of plasmid encoding SNAP-GLP-1R with a C-terminal nanoluciferase tag (SNAP-GLP-1R-Nluc) and 50 ng of fluorescent protein BRET acceptor plasmid per well of a 12-well plate, diluted with pcDNA3.1, and the assay was performed 24 hours later. SNAP-GLP-1R-Nluc was generated in *house* by PCR cloning of the nanoluciferase sequence from pcDNA3.1-ccdB-Nanoluc (a gift from Mikko Taipale; Addgene plasmid # 87067) onto the C-terminus end of the SNAP-GLP-1R vector (Cisbio), followed by site-directed mutagenesis of the GLP-1R stop codon. Cells were resuspended in HBSS containing Furimazine (1:50, Promega), seeded into 96-well half-area opaque white plates, and baseline luminescent signals recorded at 460 nm (Nluc emission peak) and 520 nm (GFP acceptor peak) or 535 nm (Venus acceptor peak) over 5 min at 37°C using a Flexstation 3 plate reader. Ligands were added, and signal was serially monitored for up to 20 min. Signal was expressed ratiometrically for each time-point as GFP or Venus acceptor divided by Nluc donor signal. The BRET ratio at each time-point was first expressed relative to the average baseline value for each well, followed by subtraction of the vehicle signal at each time-point to provide net BRET.

### 2.9 NanoBRET bystander assays

HEK293-SNAP-GLP-1R cells were transiently transfected using Lipofectamine 2000 with 50 ng of mini-G_s_-Nluc or β-arrestin-2-CyOFP [27] plus 50 ng of KRAS-Venus or Rab5-Venus (all gifts from Prof Nevin Lambert, Medical College of Georgia) per well of a 12-well plate, diluted with pcDNA3.1, and the assay was performed 24 hours later as described in Section 2.8.

### 2.10 DERET assay

The assay was performed as described previously [24]. Cells were labelled with SNAP-Lumi4-Tb (40 nM, 60 min at 37°C, in complete medium). After washing, labelled cells were resuspended in HBSS containing 24 μM fluorescein. TR-FRET signals at baseline and serially after agonist addition were recorded at 37°C using a Flexstation 3 plate reader using the following settings: λ_ex_ = 335 nm, λ_em_ = 520 and 620 nm, delay 400 μs, integration time 1500 μs. Receptor internalisation was quantified as the ratio of fluorescent signal at 620 nm to that at 520 nm, after subtraction of individual wavelength signals obtained from wells containing 24 μM fluorescein only.

### 2.11 LysoTracker TR-FRET internalisation assay

Cells were labelled with SNAP-Lumi4-Tb (40 nM, 60 minutes at 37°C, in complete medium), with LysoTracker Red DND99 (100 nM) added for the last 15 minutes of the incubation. After washing, labelled cells were resuspended in HBSS. TR-FRET signals at baseline and serially after agonist addition were recorded at 37°C using a Flexstation 3 plate reader using the following settings: λ_ex_ = 335 nm, λ_em_ = 550 and 610 nm, delay 50 μs, integration time 300 μs. Receptor translocation to acidic endosomes was quantified as the ratio of fluorescent signal at 610 nm to that at 550 nm.

### 2.12 GLP-1R clustering assay

The assay was performed as previously described [20]. Cells were dual labelled with SNAP-Lumi4-Tb (40 nM) and SNAP-Surface-649 (500 nM) for 30 min at 37°C, in complete medium. After washing, labelled cells were resuspended in HBSS. TR-FRET signals at baseline and serially after agonist addition were recorded at 37°C using a Spectramax i3x plate reader with HTRF module. GLP-1R clustering was quantified as the ratio of the fluorescence signal at 665 nm to that at 616 nm.

### 2.13 Visualisation of GLP-1R subcellular localisation

Cells were seeded onto glass coverslips and labelled with SNAP-Surface-549 (1 μM, 30 min, 37°C). After agonist stimulation and paraformaldehyde fixation, coverslips were mounted onto glass slides using Diamond Prolong antifade with DAPI. Slides were imaged using a Nikon Ti2E microscope frame with integrated hardware from Cairn Research incorporating motorised stage (ASI), LED illumination source (CoolLED) and a 100X oil immersion objective. Z-stacks were acquired and deconvolved using Deconvolutionlab2 [28] using the Richardson Lucy algorithm. Images were processed in Fiji.

### 2.14 High content microscopy trafficking assay

The assay was performed as previously described [24]. Cells were seeded into poly-D-lysine-coated, black 96-well plates. On the day of the assay, cells were labelled with BG-S-S-649 (1 μM, a gift from New England Biolabs). After washing, agonists were applied for 30 min at 37°C in complete medium. Agonists were removed, cells washed with cold HBSS and then placed on ice for subsequent steps. Mesna (100 mM in alkaline TNE buffer, pH 8.6) or alkaline TNE without Mesna was applied for 5 min, and then washed with HBSS. Microplates were then imaged using the microscope system described in Section 2.13 fitted with a 20X phase contrast objective, with data acquisition controlled by the openHCA software written for the MicroManager platform [29]. 9 random images per well were acquired for both epifluorescence and transmitted phase contrast. HBSS was then removed and replaced with fresh complete medium. Receptor was allowed to recycle for 60 min at 37°C, followed by a second Mesna application to remove any receptor that had recycled to the membrane, and the plate was re-imaged as above. Internalised SNAP-GLP-1R was quantified at both time points using Fiji as follows: 1) phase contrast images were processed using PHANTAST [30] to segment cell-containing regions from background; 2) illumination correction of fluorescence images was performed using BaSiC [31]; 3) fluorescence intensity was quantified for cell-containing regions. Agonist-mediated internalisation was determined as the mean signal for each condition normalised to signal from wells not treated with Mesna, after first subtracting non-specific fluorescence determined from wells treated with Mesna but no agonist. The percentage of reduction in residual internalised receptor after the second Mesna treatment was considered to represent recycled receptor. Recycling was then expressed as a percentage relative to the amount of receptor originally internalised in the same well.

### 2.15 Overnight stimulation surface labelling assay

INS-1-SNAP-GLP-1R cells were seeded into poly-D-lysine-coated, black 96-well plates with agonists applied in complete medium at 11 mM glucose. After an overnight incubation, agonists were removed, and cells washed with HBSS. Cells were then labelled with BG-S-S-649 (1 μM in complete medium, 30 min) before washing and imaging as in Section 2.14. Surface labelling intensity was quantified as in Section 2.14 with subtraction of signal from INS-1 832/3 cells without SNAP-GLP-1R but labelled with BG-S-S-649.

### 2.16 Insulin secretion assay

After a prior 6 hours at 3 mM glucose, INS-1 832/3 cells were seeded overnight in 96-well plates in complete medium at 11 mM glucose ± test agonists. A sample of supernatant was removed and analysed for insulin content by HTRF (Wide Range Insulin HTRF Kit, Cisbio). Results were expressed relative to insulin released with 11 mM glucose alone.

### 2.17 In vivo study

Animals were maintained in specific pathogen-free facilities, with *ad lib* access to food (except prior to fasting studies) and water. Studies were regulated by the UK Animals (Scientific Procedures) Act 1986 of the U.K. and approved by Imperial College London (Project License PB7CFFE7A). Male C57Bl/6 mice (8-10 weeks of age, weight 25 - 30g, supplied by Charles River, UK) were injected intra-peritoneally (IP) during the light phase with 2.4 nmol/kg agonist prepared in 100 μl 0.9% NaCl, or vehicle. Food was removed from the cages at this point until the end of the study. After an 8-hour delay, mice were injected IP with 20% dextrose (2 g/kg). Blood glucose levels were monitored immediately before glucose administration and at 20-min intervals thereafter from tail vein blood samples using a hand-held glucose meter.

### 2.18 Data analysis

All analyses were performed using Prism 8 (GraphPad Software). The average of within-assay replicates was counted as one biological replicate. For equilibrium binding studies, Luxendin645 and exendin-4-Cy5 K_d_ values were fitted using a “one site – specific binding” model. Test ligand K_i_ values were determined using a “one site – fit K_i_” model. Functional concentration response data were fitted using 3-parameter logistic fitting, with constraints applied where appropriate. Bias analysis was performed by the calculation of transduction coefficients (i.e. τ/K_A_ values) as previously described [9,32,33], with subtraction of the Log(τ/K_A_) value for one pathway (cAMP or mini-G_s_) from the second pathway (β-arrestin-2) to give ΔLog(τ/K_A_). The assays for which bias was calculated were performed with both pathways assessed in parallel using the same ligand stock, allowing statistical comparisons on a per-assay basis. In the instances when a composite parameter was determined from two assay types not performed in parallel (i.e. the pEC_50_ - pK_i_ measurement in Figure 1), error propagation was performed. Statistical comparisons were performed by t-test, one-way ANOVA or two-way ANOVA as appropriate. Paired or matched analyses were performed where permitted by the experimental design. Specific post-hoc tests are indicated in the legends. Statistical significance was assigned if p<0.05. Data are represented as mean ± standard error of the mean (SEM) throughout, with individual replicates indicated where possible.

**Figure 1.**
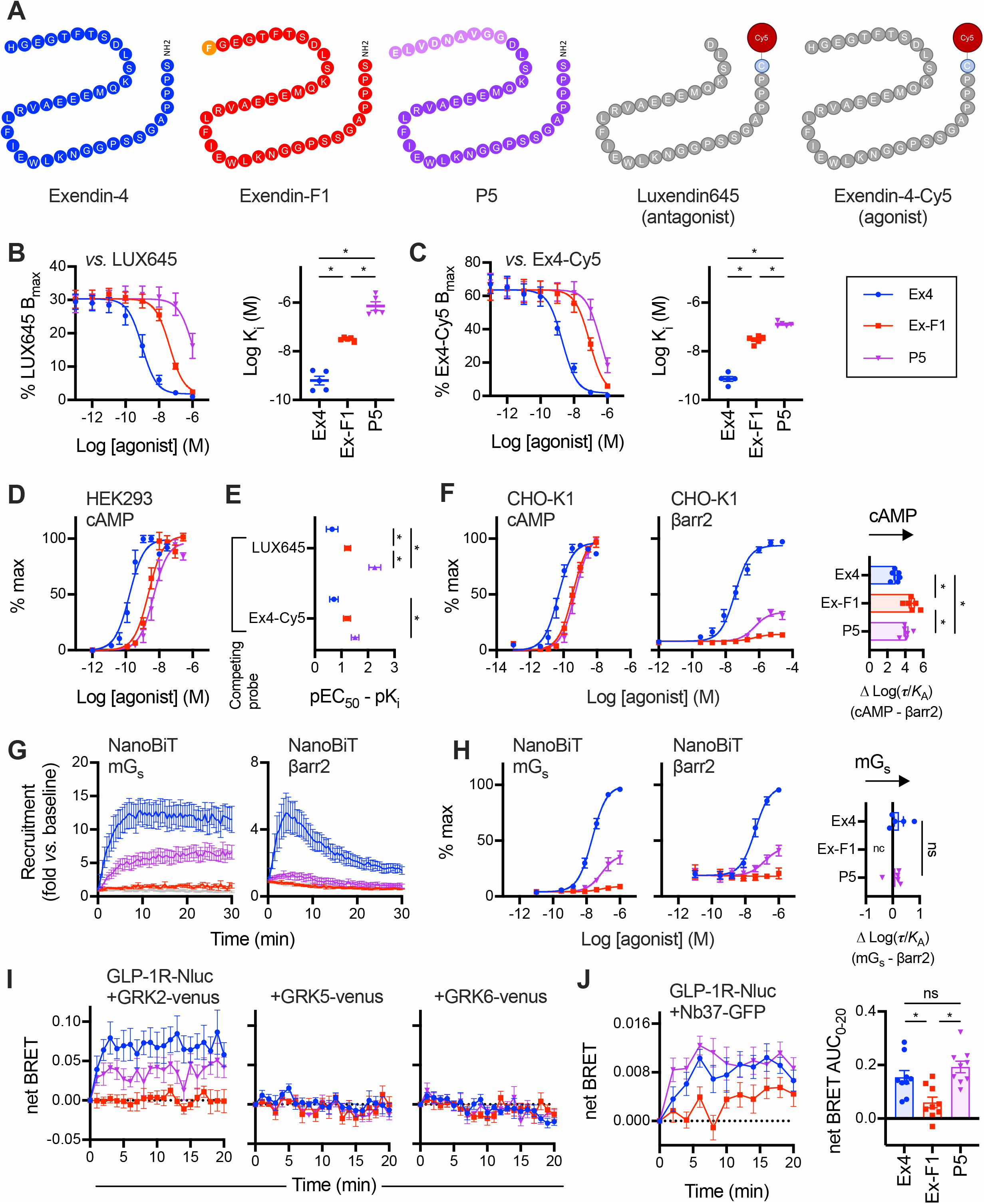
Pharmacological characterisation of GLP-1R agonists. (**A**) Schematic showing amino acid sequences of peptide ligands used in this study in single letter code. (**B**) Competitive binding of each ligand in competition with 10 nM Luxendin645 (LUX645), *n*=5, with calculated K_i_ shown and compared by one-way randomised block ANOVA with Tukey’s test. (**C**) As for (B) but with 5 nM exendin-4-Cy5 (Ex4-Cy5) as the competing ligand, *n*=5. (**D**) cAMP production in HEK293-SNAP-GLP-1R cells, 30-min stimulation, normalised to global maximum response, with 3-parameter fit shown, *n*=5. (**E**) cAMP potency from (D) expressed relative to affinity from (B) and (C), with error propagation, and comparison by two-way ANOVA with Tukey’s test. (**F**) cAMP production and recruitment of β-arrestin-2 (βarr2) in PathHunter CHO-K1-βarr2-EA cells, 30-min stimulation, normalised to full agonist global maximum response, with 3-parameter fit, *n*=6, and bias determination as ΔLog(τ/K_A_) (cAMP *versus* β-arrestin-2 for each ligand) shown and compared by one-way randomised block ANOVA with Tukey’s test. (**G**) 100 nM ligand-induced recruitment of mini-G_s_ (mG_s_) and β-arrestin-2 to GLP-1R, measured by nanoBiT complementation in transiently transfected HEK293 cells, *n*=5. (**H**) Ligand-induced recruitment of mini-G_s_ (mG_s_) and β-arrestin-2 to GLP-1R at 5 min, measured by nanoBiT complementation in transiently transfected HEK293 cells, *n*=5, with bias determination as ΔLog(τ/K_A_) shown and compared by paired t-test (exendin-4 *versus* P5). (**I**) Recruitment of GRK2-Venus, GRK5-Venus, or GRK6-Venus to GLP-1R-Nluc in transiently transfected HEK293T cells, with 100 nM agonist applied, all *n*=6. (**J**) Recruitment of Nb37-GFP to GLP-1R in response to 100 nM agonist in HEK293T cells, *n*=9, with AUC shown and compared by one-way randomised block ANOVA with Tukey’s test. * p<0.05 by indicated statistical test. Data are represented as mean ± SEM, with individual replicates shown where possible.

## 3 Results

### 3.1 Contrasting efficacy and affinity with P5 *versus* exendin-F1

We first measured the equilibrium binding affinities of exendin-4, exendin-F1 and P5 (Figure 1A) in HEK293 cells stably expressing SNAP-GLP-1R [34], using a competitive time-resolved FRET assay in which the receptor N-terminus is labelled with the energy donor Lumi4-Tb to detect binding of Cy5-conjugated GLP-1R antagonist exendin(9-39) (“Luxendin645”) [19] or the equivalent agonist, exendin-4-Cy5 [18], applied in competition with unlabelled test peptides (Figure 1B and 1C, Table 1). Both G protein-biased ligands displayed significantly lower affinity for receptor binding than exendin-4; of note, the K_i_ for P5 was 22-fold higher than for exendin-F1 when measured using Luxendin645 as the competing probe but only 4-fold higher with exendin-4-Cy5. Measurement of cAMP production in the same cell model showed reduced potency for both exendin-F1 and P5 (Figure 1D, Table 1), but the differences between agonists were smaller than for equilibrium binding affinity, suggesting ligand-specific differences in coupling of receptor occupancy to cAMP production (Figure 1E). This observation has been made before for P5 [35] and exendin-F1 [34], but here the ligands are compared directly for the first time.

**Table 1.** Parameter estimates for pharmacological responses to GLP-1R agonists. Mean ± SEM for 3-parameter fit-derived potency and efficacy estimates from Figures 1 and 2. For E_max_, where all ligands were full agonists, values are normalised to a “global” E_max_ obtained for each assay, whereas where only exendin-4 was a full agonist, exendin-F1 and P5 E_max_ values are expressed relative to exendin-4. Statistical comparisons are by one-way randomised block ANOVA with Tukey’s test. Statistical testing for E_max_ values was performed on data prior to normalisation. * p<0.05 *versus* exendin-4; ^#^ p<0.05 exendin-F1 *versus* P5. n.c. = not calculable.

| Assay and cell model | Exendin-4 |  | Exendin-F1 |  | P5 |  |
| --- | --- | --- | --- | --- | --- | --- |
|  | pEC<sub>50</sub><br>(M) | E<sub>max</sub><br>(%) | pEC<sub>50</sub><br>(M) | E<sub>max</sub><br>(%) | pEC<sub>50</sub><br>(M) | E<sub>max</sub><br>(%) |
| cAMP (HEK293-SNAP-GLP-1R) | 9.9 ± 0.1 | 95 ± 5 | 8.7 ± 0.1 * | 98 ± 6 | 8.4 ± 0.1 *, # | 92 ± 5 |
| cAMP (PathHunter) | 10.3 ± 0.1 | 97 ± 1 | 9.4 ± 0.1 * | 105 ± 3 | 9.3 ± 0.1 * | 107 ± 2 * |
| β-arrestin2 (PathHunter) | 7.5 ± 0.2 | 100 | 6.4 ± 0.2 * | 15 ± 0 * | 6.3 ± 0 * | 36 ± 3 *, # |
| NanoBiT mini-G<sub>s</sub> (HEK293T) | 7.7 ± 0.1 | 100 | 7.1 ± 0.3 | 10 ± 2 * | 7.1 ± 0.1 | 38 ± 7 * |
| NanoBiT β-arrestin2 (HEK293T) | 7.4 ± 0.1 | 100 | n.c. | n.c. | 6.8 ± 0.2 * | 46 ± 7 * |
| Internalisation by microscopy (HEK293-SNAP-GLP-1R) | 8.3 ± 0.1 | 93 ± 8 | 7.2 ± 0.1 * | 17 ± 4 * | 6.8 ± 0.2 * | 64 ± 5 *, # |
| Internalisation by DERET (HEK293-SNAP-GLP-1R) | 8.5 ± 0.2 | 100 | n.c. | n.c. | n.c. | n.c. |
| Lysosomal redistribution (HEK293-SNAP-GLP-1R) | 8.0 ± 0.2 | 100 | n.c. | n.c. | n.c. | n.c. |

Balance between G protein pathway engagement and β-arrestin-2 recruitment was assessed using two approaches. Firstly, using the PathHunter system [9], cAMP production and β-arrestin-2 recruitment were measured in parallel, which highlighted how exendin-F1 and P5 show low efficacy as well as reduced potency for β-arrestin-2 (Figure 1F, Table 1). β-arrestin-2 recruitment efficacy was greater for P5 than for exendin-F1. Comparison of transduction ratios (T/*K*_A_) determined using the operational model for each pathway [33] indicated a substantial degree of bias in favour of cAMP production for both P5 and exendin-F1, with the effect being most marked for the latter (~60-fold *versus* 15-fold). Secondly, recruitment of mini-G_s_ and β-arrestin-2 to GLP-1R were measured by nanoBiT complementation [25,36], with response kinetics shown in Figure 1G and concentration-responses at 5 min in Figure 1H (see also Table 1). These analyses confirmed low efficacy β-arrestin-2 recruitment but also reduced recruitment efficacy for mini-G_s_ with exendin-F1 and P5. Maximum responses with exendin-F1 were reduced in comparison to P5 in both pathways, to the extent that the β-arrestin-2 response was not amenable to logistic curve fitting for the former ligand. Interestingly, no significant bias could be detected for P5 *versus* exendin-4 from these data; bias for exendin-F1 could not be determined due to the lack of a quantifiable β-arrestin-2 response.

We also investigated the relative propensity for each ligand to recruit G protein-coupled receptor kinases (GRKs) to the GLP-1R, an intermediate step that typically precedes recruitment of β-arrestins to GPCRs, including GLP-1R [37,38]. Even at a high (100 nM) stimulatory concentration, exendin-F1 showed very minimal recruitment of GRK2 to GLP-1R compared to exendin-4, as measured by BRET between SNAP-GLP-1R tagged at the C-terminus with nanoluciferase (SNAP-GLP-1R-Nluc) and GRK2-Venus, with an intermediate effect for P5 (Figure 1I). Ligand-induced changes were barely detectable for GRK5 and 6.

We also designed a BRET-based sensor strategy to monitor differences in ligand-induced activation (as opposed to recruitment) of endogenous G proteins by co-expressing SNAP-GLP-1R-Nluc with GFP-tagged nanobody-37 (Nb37), a genetically encoded intrabody that recognises the active conformation of Gα_S_ [26]. Interestingly, the Nb37 response to exendin-F1, but not P5, was reduced in comparison to exendin-4 in HEK293 cells (Figure 1J). The low dynamic range of this sensor configuration precluded concentration-response analyses.

Overall, these data highlight how exendin-F1 has a higher GLP-1R binding affinity than P5 but lower efficacy for recruitment of mini-G_s_, GRK2 and β-arrestin-2, as well as reduced Gα_s_ activation, which implies greater coupling efficiency of GLP-1R occupancy to intracellular responses with P5 *versus* exendin-F1. However, the higher affinity of exendin-F1 results in comparable acute cAMP signalling responses between both biased agonists in the heterologous cell system used for these studies.

### 3.2 GLP-1R trafficking with exendin-F1 and P5

An altered membrane trafficking profile characterised by reduced endocytosis and faster recycling is thought to be an important component of the action of biased GLP-1R agonists such as exendin-F1 [9], but has not been tested for P5. Using high content microscopy [24], we confirmed that both exendin-F1 and P5 showed reduced GLP-1R internalisation propensity but faster recycling than exendin-4 (Figure 2A, Table 1). The reduction in efficacy was again most pronounced with exendin-F1 but potency was slightly lower with P5; relative potencies for each ligand *versus* exendin-4 were in line with those for cAMP production in the same cell model. Higher resolution images of GLP-1R subcellular localisation with each ligand are shown in Figure 2B, revealing that a substantial proportion of GLP-1R remains at the plasma membrane after stimulation with both biased GLP-1R agonists, in comparison to exendin-4. Kinetics of GLP-1R internalisation were assessed by diffusion-enhanced resonance energy transfer (DERET) [39], with very little response detected with either exendin-F1 or P5 below 100 nM (Figure 2C, Table 1).

**Figure 2.**
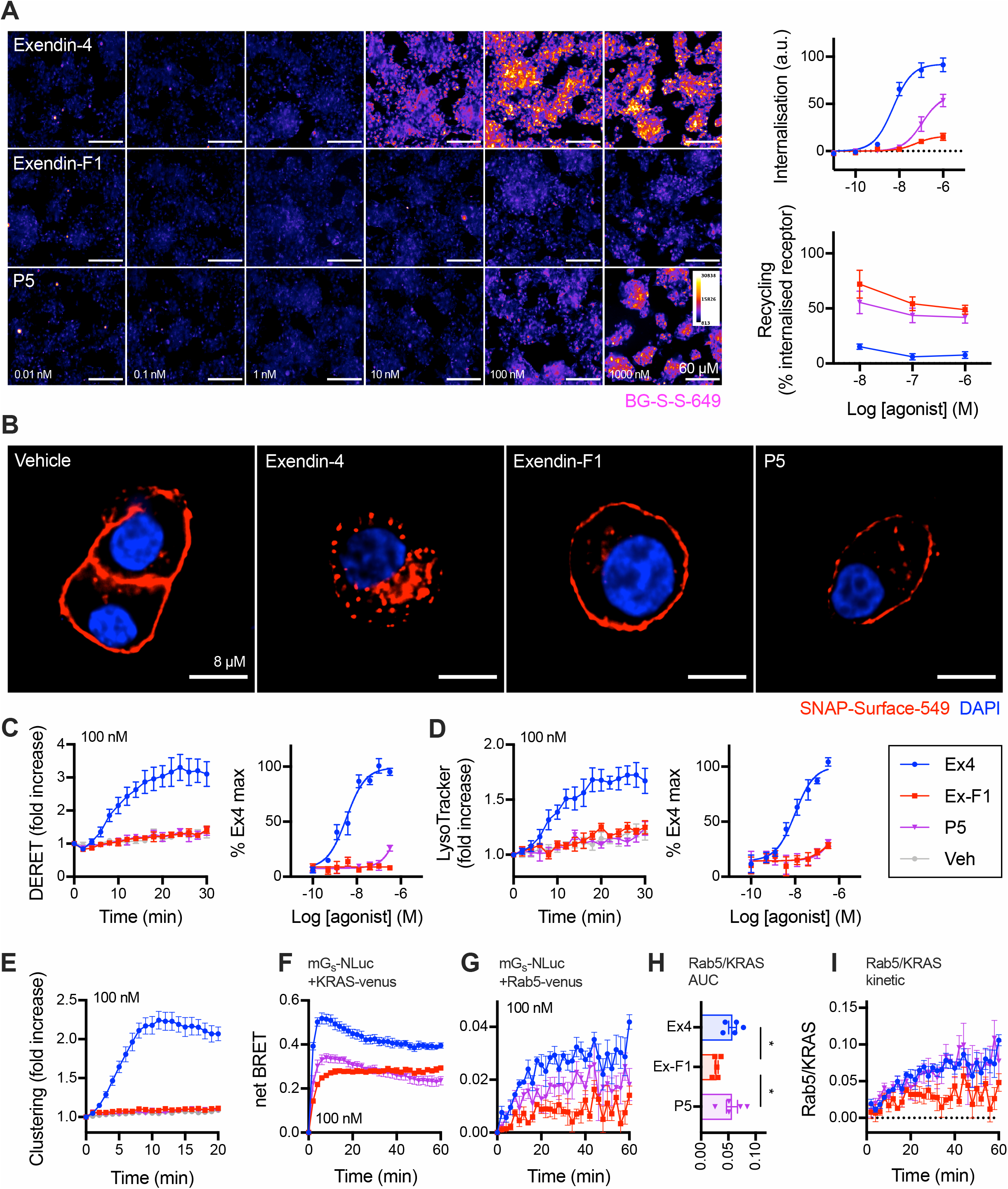
Trafficking responses of biased GLP-1R agonists. (**A**) GLP-1R internalisation (30 min) and recycling (60 min) with each GLP-1R agonist in HEK293-SNAP-GLP-1R cells. Representative cropped images are shown for the internalisation step, with quantification from *n*=5 experiments. Scale bar = 60 μm. (**B**) Representative high-resolution images showing GLP-1R internalisation in HEK293-SNAP-GLP-1R cells with 100 nM ligand, 30 min. Scale bar = 8 μm. (**C**) GLP-1R internalisation in HEK293-SNAP-GLP-1R cells measured by DERET, with kinetic response for 100 nM agonist shown and quantification from 30-min AUC also indicated, *n*=5. (**D**) As for (C) but for GLP-1R trafficking to lysosomal compartment. (**E**) GLP-1R clustering in HEK293-SNAP-GLP-1R cells stimulated with 100 nM agonist, *n*=5. (**F**) Recruitment of mini-G_s_-Nluc to plasma membrane (KRAS-Venus marker) in HEK293-SNAP-GLP-1R cells stimulated with 100 nM agonist, *n*=5. (**G**) As for (F) but recruitment to early endosomes (Rab5-Venus marker). (**H**) AUC ratio indicating balance of Rab5 to KRAS mini-G_s_-Nluc BRET signals from (F) and (G), with statistical comparison by one-way randomised block ANOVA with Tukey’s test. (**I**) Ratio of BRET signals from (F) and (G) expressed at each time-point. * p<0.05 by indicated statistical test. Data are represented as mean ± SEM, with individual replicates shown where possible.

We also developed a time-resolved FRET (TR-FRET) assay to monitor translocation of lanthanide-labelled SNAP-GLP-1R to the late endolysosomal compartment, which was labelled using the chemical endolysomotropic dye LysoTracker DND99. This highlighted how exendin-4 rapidly targets internalised GLP-1Rs towards this degradative compartment (Figure 2D, Table 1). In these studies, performed in parallel to DERET measurements, the EC_50_ for exendin-4-induced GLP-1R lysosomal localisation was somewhat higher than for internalisation *per se*, implying that not all internalised GLP-1Rs are targeted to the lysosome.

As spatial reorganisation of activated GLP-1Rs into tightly constrained nanodomains precedes endocytosis [20], we also compared GLP-1R clustering with each ligand using a dual-labelling TR-FRET assay. GLP-1R clustering was barely detectable at 100 nM agonist with exendin-F1 or P5, but exendin-4 produced a robust response (Figure 2E).

Agonist-internalised GLP-1Rs continue to generate cAMP signals from the endosomal compartment [40], with this process being ligand specific [41]. To monitor the distribution of GLP-1Rs in their active state between plasma membrane and endosomes with exendin-4, P5 and exendin-phe1, we co-expressed mini-G_s_ tagged with nanoluciferase with Venus-tagged markers of the plasma membrane (KRAS) or early endosome (Rab5) [25]. In this bystander configuration, the location of GLP-1R in its active conformation is inferred when mini-G_s_ is recruited to the vicinity of the relevant compartment marker leading to an increase in BRET signal. In stable HEK293-SNAP-GLP-1R cells, 100 nM exendin-4 induced robust and rapid translocation of mini-G_s_ to the plasma membrane, followed by a gradual decline; a similar pattern, but with a lower peak, was seen with P5 (Figure 2F). For exendin-F1, the peak response was further reduced, but did not fall below the peak level for the full 30-minute stimulation period. Mini-G_s_ recruitment to Rab5-positive early endosomes was of slower onset than at the plasma membrane, and response magnitude showed a rank order matching that of GLP-1R endocytosis (exendin-4 > P5 > exendin-F1; Figure 2G). Expressing ligand-specific Rab5 and KRAS signals ratiometrically from their AUC (Figure 2H), or over time (Figure 2I), suggested P5 engenders a similar balance between endosomal and plasma membrane activity to exendin-4, but exendin-F1 delivers a predominantly plasma membrane delimited response. We did not attempt to perform full concentration responses with these assays due to the low dynamic range of the Rab5 BRET assay.

Therefore, both P5 and exendin-F1 show reduced GLP-1R endocytosis and accelerated recycling, with this effect being most dramatic for exendin-F1. The differences in GLP-1R internalisation between P5 and exendin-F1 are mirrored by their respective tendencies to elicit GLP-1R activity at early endosomes.

### 3.3 Beta cell and *in vivo* effects of exendin-F1 *versus* P5

GLP-1R-specific signalling was confirmed for each peptide by comparing acute cAMP responses in the pancreatic beta cell line INS-1 832/3 [21] and a CRISPR/Cas9-derived subclone thereof lacking endogenous GLP-1R or GIPR expression [22] (Figure 3A, Table 2). Interestingly, efficacy with P5 was slightly higher than with the other two ligands in GLP-1R-expressing cells.

**Figure 3.**
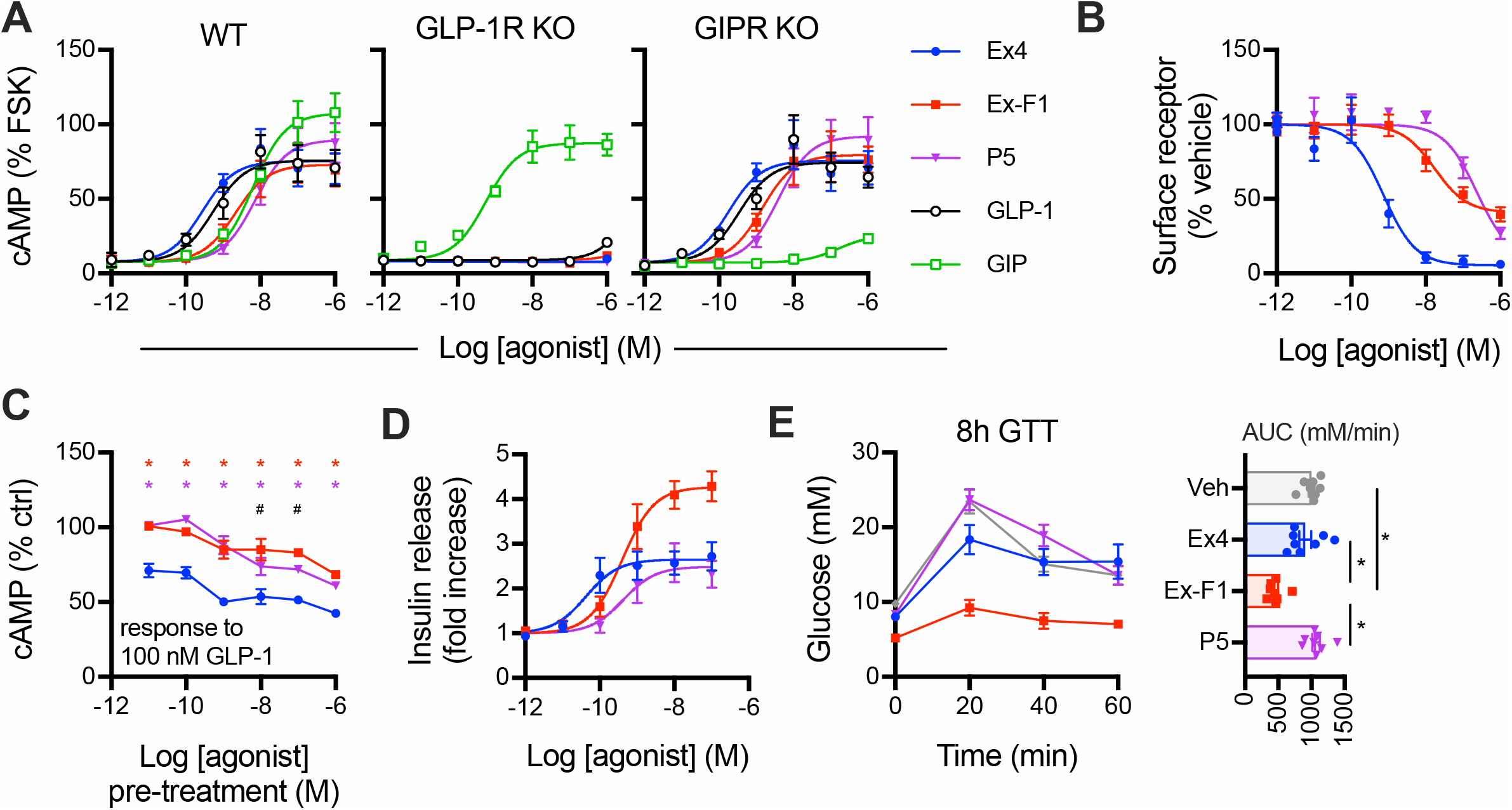
Responses in beta cells and anti-hyperglycaemic efficacy. (**A**) Acute cAMP signalling in INS-1 832/3 cells with and without endogenous GLP-1R or GIPR, as indicated, for 10-min stimulation with 500 μM IBMX, expressed relative to forskolin (FSK; 10 μM) response, 3-parameter fits shown, *n*=4. (**B**) Residual surface SNAP-GLP-1R after overnight treatment of INS-1 832/3 cells with indicated agonist concentrations, 3-parameter fits shown, *n*=5. (**C**) Homologous desensitisation assay in INS-1 832/3 cells treated overnight with indicated agonist concentration, followed by a 1-hour recovery period and stimulation with 100 nM GLP-1 plus 500 μM IBMX, *n*=4. Responses are expressed relative to vehicle pre-treated cells, with two-way randomised block ANOVA with Sidak’s test performed. (**D**) Cumulative insulin secretion in INS-1 832/3 cells treated with indicated agonist overnight at 11 mM glucose, expressed as a fold change relative to response to zero agonist condition, *n*=5. (**E**) Intraperitoneal glucose tolerance test (2 g/kg glucose) performed in lean male C57Bl/6 mice, 8 hours after administration of 2.4 nmol/kg agonist, *n*=8 per group, with AUC comparisons by one-way ANOVA with Tukey’s test. * p<0.05 by indicated statistical test; for (C), red asterisk indicates exendin-F1 *versus* exendin-4, purple asterisk indicates P5 *versus* exendin-4, # indicates exendin-F1 *versus* P5. Data are represented as mean ± SEM, with individual replicates shown where possible.

**Table 2.** Parameter estimates for responses in beta cell models. Mean ± SEM for potency and efficacy estimates from Figure 3. E_max_ values are reported as in the figure, i.e. as a % of forskolin response for INS-1 cAMP results, % remaining surface receptor for GLP-1R downregulation assay, and as fold change response *versus* 11 mM glucose for the insulin secretion assay. Statistical comparisons are by one-way randomised block ANOVA with Tukey’s test. * p<0.05 *versus* exendin-4; ^#^ p<0.05 exendin-F1 *versus* P5. n.c. = not calculable.

| Assay and cell model | Exendin-4 |  | Exendin-F1 |  | P5 |  |
| --- | --- | --- | --- | --- | --- | --- |
|  | pEC<sub>50</sub><br>(M) | E<sub>max</sub> | pEC<sub>50</sub><br>(M) | E<sub>max</sub> | pEC<sub>50</sub><br>(M) | E<sub>max</sub> |
| cAMP (INS-1 832/3, wild-type) | 9.6 ± 0.1 | 76 ± 11 | 8.6 ± 0.2 * | 74 ± 12 | 8.1 ± 0.2 *, # | 91 ± 13 *, # |
| cAMP (INS-1 832/3, GLP-1R KO) | n.c. | n.c. | n.c. | n.c. | n.c. | n.c. |
| cAMP (INS-1 832/3, GIPR KO) | 9.8 ± 0.1 | 76 ± 13 | 8.8 ± 0.2 * | 80 ± 14 | 8.3 ± 0.2 *, # | 93 ± 15 *, # |
| GLP-1R downregulation (INS-1-SNAP-GLP-1R) | 9.2 ± 0.2 | 6 ± 2 | 7.7 ± 0.2 * | 39 ± 5 * | 6.6 ± 0.2 *, # | 6 ± 3 # |
| Insulin secretion (INS-1 832/3) | 10.3 ± 0.1 | 2.7 ± 0.3 | 9.4 ± 0.1 * | 4.3 ± 0.3 * | 9.1 ± 0.3 * | 2.5 ± 0.4 # |

Previous work has demonstrated how prolonged stimulation with GLP-1R agonists with different trafficking phenotypes leads to variable levels of receptor downregulation, which can influence their insulinotropic potential [9,10]. In SNAP-GLP-1R stably expressed in INS-1 832/3 cells lacking endogenous GLP-1R, referred to as INS-1-SNAP-GLP-1R cells [42], both exendin-F1 and P5 showed markedly increased preservation of surface GLP-1R levels compared to exendin-4 (Figure 3B, Table 2). Exendin-F1 led to loss of surface GLP-1R at lower concentrations than P5, as expected from its higher affinity, but the maximum loss of surface receptor effect was less marked.

To determine the functional impact of differential GLP-1R downregulation, we measured cAMP responses to a fixed concentration of GLP-1 after a prior 16-hour pre-treatment phase with each exendin analogue. Both biased ligands produced substantially less homologous desensitisation than exendin-4 (Figure 3C). Whilst concentration responses could not be accurately quantified by logistic curve fitting of data from assay repeats, P5 showed greater desensitisation than exendin-F1 at pre-treatment concentrations upwards of 10 nM.

We also measured cumulative insulin secretion with each ligand after an overnight stimulation as a therapeutically relevant readout of sustained pharmacological agonism in beta cells. Here, there was a stark difference in insulinotropic efficacy, with exendin-F1 treatment yielding approximately twice as much insulin secretion as exendin-4 and P5 (Figure 3D, Table 2). Moreover, assessment of anti-hyperglycaemic efficacy in mice by intraperitoneal glucose tolerance testing, performed 8 hours after agonist administration to allow desensitisation-related effects to emerge as previously performed [9,10], showed a pattern compatible with apparent greater sustained action with exendin-F1 (Figure 3E). Note that both P5 and exendin-F1 have been demonstrated to show identical pharmacokinetics to exendin-4 [8,9].

## 4 Discussion

In this study we have compared two exendin-4 analogues, P5 and exendin-F1, that were the first two synthetic, orthosteric GLP-1RAs reported to show biased agonism favouring G protein-dependent signalling over β-arrestin recruitment and tested *in vivo* for their metabolic effects. Biased GLP-1R agonism has emerged as a promising therapeutic strategy for T2D on the basis of preclinical evaluations [8,9,11] and the recent recognition that some of the beneficial effects of the dual GLP-1R/GIPR agonist tirzepatide may be due to biased agonism at the GLP-1R [14]. Whilst in our study we confirmed the exendin-F1 and P5 do indeed show selective reduction in β-arrestin recruitment, their patterns of engagement with intracellular effectors, trafficking profiles, subcellular signalling localisation and insulinotropic efficacy were in fact rather different.

It is notable that both exendin-F1 and P5, considered G protein-biased from previous reports [8,9], are in fact low efficacy agonists for mini-G_s_ engagement compared to the full agonist exendin-4. We verified bias in favour of cAMP production over β-arrestin-2 recruitment in the PathHunter system, but interestingly, the same operational analysis did not indicate bias between mini-G_s_ and β-arrestin-2 recruitment for P5 (while for exendin-F1 the β-arrestin-2 response was undetectable, precluding formal analysis). There is increasing recognition that bias comparisons between readouts encompassing significant signal amplification (e.g. cAMP) and those without amplification (e.g. β-arrestin recruitment) are susceptible to system non-linearity that may confound current models [43,44]. However, if seen from the pragmatic point of view that efficacy is a key driver of the manifestations of biased agonism [43], considering P5 and exendin-F1 as G protein-biased appear appropriate, as in both cases β-arrestin-2 recruitment is even more markedly reduced than mini-G_s_ recruitment.

Our whole cell binding assays indicated lower affinity for P5 than for exendin-F1. Indeed, P5 was previously identified as a low affinity agonist, with a reported ~100-fold lower affinity than GLP-1 in competition with iodinated exendin(9-39) [35]. Our P5 pK_i_ measurements were dependent on the choice of competing fluorescent probe, with apparently lower affinity measured when the antagonist LUXendin645 was used rather than the equivalent agonist exendin-4-Cy5. In a previous report, the absence of G proteins, or expression of a dominant negative Gα_s_, led to smaller differences between P5 and exendin-4 affinity in insect cells [35]. Thus, probe-dependency of measured pK_i_ with P5 could represent a higher affinity GLP-1R active-state complex triggered by exendin-4-Cy5-induced coupling to Gα_s_ that is not seen with LUXendin645 [45]. Additionally, whilst our whole cell binding assays were performed at low temperature and with metabolic inhibitors to avoid the effects of ATP-dependent GLP-1R redistribution and endocytosis on binding phenomena, we cannot absolutely exclude these as ligand-specific confounders. Affinity measurements in cell-free reconstituted systems could be performed to address this issue.

Contrasting with its higher affinity, maximum responses for multiple readouts were further reduced for exendin-F1 than for P5. Thus, recruitment of mini-G_s_, GRK2 and β-arrestin-2 to the receptor, and activation of Gα_s_ as measured by Nb37 recruitment to SNAP-GLP-1R-Nluc, were measurably lower with high doses of exendin-F1 *versus* P5, as was GLP-1R endocytosis. The counterbalancing of effects on affinity *versus* efficacy appears to result in similar acute cAMP potency estimates for both ligands (albeit slightly reduced with P5, depending on the cell system), with P5 relying on greater coupling to intracellular effectors in the face of lower affinity. This distinction may be important as efficacy- *versus* affinity-driven signalling can manifest differently in tissues with greater or lesser sensitivity to GLP-1R stimulation, due to factors such as expression levels of GLP-1R, signalling intermediates and downstream effectors [46]. Specifically, in the presence of adequate GLP-1R expression, the low acute efficacy for both G protein and β-arrestin engagement of exendin-F1 is still sufficient to fully activate cAMP/PKA signalling either due to either signal amplification or differences in capacity to induce activation of recruited G proteins, whilst at the same time avoiding β-arrestin-mediated desensitisation and downregulation phenomena that otherwise limit response duration. In contrast, the same ligand might fail to induce high amplitude responses in tissues with lower GLP-1R expression. This tissue selectivity could potentially influence the risk of side effects when used therapeutically.

After endocytosis, a number of GPCRs are known to continue to signal from the endosomal compartment [47]. Pharmacological and genetic inhibition of GLP-1R endocytosis attenuates cAMP production [48,49], and Gα_s_ was found to colocalise with GLP-1R in early endosomes [40]. In our study, we have monitored ligand-specific patterns of redistribution of GLP-1R in its Gα_s_-preferring active conformation from the plasma membrane to early endosomes. Here, P5 led to more active GLP-1Rs at Rab5-positive early endosomes than exendin-F1, in line with the greater total GLP-1R internalisation recorded with this ligand. This could be important given that “location bias” can modulate agonist effects, as the same intracellular signalling events originating at the endosome *versus* plasma membrane may result in distinct downstream actions [50]. Endosomal signalling is often referred to as being responsible for sustained responses [51]. However, our results call into question the relative importance of this phenomenon in controlling the duration of action of GLP-1RAs in the therapeutic setting, as exendin-F1 showed the least tendency to promote GLP-1R activation at the endosomal compartment but the greatest maximal insulin secretion after prolonged incubation with beta cells, and a greater glucose-lowering effect than P5 in a delayed glucose tolerance test in mice. This is presumably attributable to a sufficiently enhanced avoidance of receptor desensitisation and downregulation, both of which ultimately limit the global capacity for signalling for the GLP-1R, to the point that any potential advantages of endosomal signalling no longer dominate. Ultimately, the balance between the positive and negative contributions of all the above-mentioned factors is likely to determine the overall capacity of each pharmacological agonist for sustained action.

Despite its reduced importance in a pharmacological setting, endosomal signalling is likely to play a more prominent role in mediating physiological GLP-1R effects, as the lower plasma concentrations and short circulatory half-life of GLP-1 and other proglucagon-derived peptides means that the fraction of downregulated GLP-1R is likely to be minimal. Moreover, it should be emphasised that our mini-G_s_ bystander BRET assay does not measure endosomal signalling *per se,* but the presence of activated GLP-1R at particular locations. Targeted biosensors to monitor cAMP generation [41] or PKA activation [52] are available, and Nb37 redistribution can be observed by microscopy [26], although these tend to be lower throughput methods.

At the outset of the present study, we anticipated that P5 would in fact be highly insulinotropic when assessed using the same methodology [9] used to reveal the impact of GLP-1R desensitisation and time-dependent advantages of exendin-F1, but this proved not to be the case. It appears that P5 achieves a less favourable balance between “beneficial” signalling and “non-beneficial” desensitisation which ultimately limits the amplitude of the sustained insulin release response, which was in fact similar to that of exendin-4. The potent effects of P5 on blood glucose lowering reported by Zhang *et al* [8] were observed in the “hyper-acute” setting, i.e. with a different approach to the delayed IPGTT used in our study, as well as being corroborated by improvements in HbA1c after chronic administration. Our results do not shed any light on how P5 obtains its pronounced anti-hyperglycaemic effects, which remain hard to explain as P5 was less insulinotropic than exendin-4 and had no differential effect on insulin sensitivity. An insulin-independent mechanism remains a possibility but was not explored in the study of Zhang *et al* [8]. Chronic treatment with P5 led to various metabolic changes that exceed those of the same dose of exendin-4, including increased adipose tissue hyperplasia and reduced inflammation, along with increased circulating GIP and reduced circulating resistin [8]. These may be relevant in the chronic setting but unlikely to contribute to the hyperacute effects of P5 on glucose levels.

In summary, whilst P5 and exendin-F1 superficially resemble each other at the pharmacological level, our study highlights that emphasising lower acute efficacy for transducer coupling, but higher affinity or potency, may be a more viable strategy to achieve therapeutically optimised GLP-1R biased agonism.

## Acknowledgements

The Section of Endocrinology and Investigative Medicine is funded by grants from the MRC, BBSRC, NIHR, and is supported by the NIHR Biomedical Research Centre Funding Scheme. The views expressed are those of the authors and not necessarily those of the funders. This project was supported by an MRC project grant (MR/R010676/1) to A.T., B.J., S.R.B., and G.A.R. The European Federation for the Study of Diabetes has supported B.J. and A.T. in this work. A.T. also acknowledges funding from Diabetes UK. B.J. acknowledges support from the Academy of Medical Sciences, Society for Endocrinology, The British Society for Neuroendocrinology, the European Federation for the Study of Diabetes, and an EPSRC capital award. D.J.H. was supported by MRC (MR/N00275X/1 and MR/S025618/1) and Diabetes UK (17/0005681) Project Grants. This project has received funding from the European Research Council (ERC) under the European Union’s Horizon 2020 research and innovation programme (Starting Grant 715884 to D.J.H.). G.A.R. was supported by Wellcome Trust Investigator (212625/Z/18/Z) Awards, MRC Programme (MR/R022259/1) and Experimental Challenge Grant (DIVA, MR/L02036X/1), and Diabetes UK (BDA/11/0004210, BDA/15/0005275, BDA 16/0005485) grants. This work has received support from the EU/EFPIA/Innovative Medicines Initiative 2 Joint Undertaking (RHAPSODY grant No 115881) to G.A.R. V.S. was supported by a Diabetes UK Harry Keen Fellowship.

## Author contributions

Amaara Marzook: Investigation, Formal analysis, Writing – Review and Editing, Visualization

Shiqian Chen: Investigation, Formal analysis, Writing – Review and Editing

Phil Pickford: Investigation, Formal analysis, Writing – Review and Editing

Maria Lucey: Investigation, Formal analysis, Writing – Review and Editing

Yifan Wang: Investigation, Writing – Review and Editing

Ivan R Corrêa, Jr: Resources, Writing – Review and Editing

Johannes Broichhagen: Resources, Writing – Review and Editing

David J Hodson: Resources, Writing – Review and Editing

Victoria Salem: Supervision, Writing – Review and Editing

Guy A Rutter: Funding acquisition, Writing – Review and Editing

Tricia M Tan: Supervision, Writing – Review and Editing

Stephen R Bloom: Funding acquisition, Writing – Review and Editing

Alejandra Tomas: Conceptualization, Funding acquisition, Writing – Review and Editing

Ben Jones: Conceptualization, Funding acquisition, Investigation, Formal analysis, Writing – Original Draft, Visualization, Project administration.

